# Tuning Spatial Profiles of Selection Pressure to Modulate the Evolution of Resistance

**DOI:** 10.1101/230854

**Authors:** Maxwell G. De Jong, Kevin B. Wood

## Abstract

Spatial heterogeneity plays an important role in the evolution of drug resistance. While recent studies have indicated that spatial gradients of selection pressure can accelerate resistance evolution, much less is known about evolution in more complex spatial profiles. Here we use a stochastic toy model of drug resistance to investigate how different spatial profiles of selection pressure impact the time to fixation of a resistant allele. Using mean first passage time calculations, we show that spatial heterogeneity accelerates resistance evolution when the rate of spatial migration is sufficiently large relative to mutation but slows fixation for small migration rates. Interestingly, there exists an intermediate regime—characterized by comparable rates of migration and mutation—in which the rate of fixation can be either accelerated or decelerated depending on the spatial profile, even when spatially averaged selection pressure remains constant. Finally, we demonstrate that optimal tuning of the spatial profile can dramatically slow the spread and fixation of resistant subpopulations, even in the absence of a fitness cost for resistance. Our results may lay the groundwork for optimized, spatially-resolved drug dosing strategies for mitigating the effects of drug resistance.

---

Drug resistance is a rapidly growing public health threat and a central impediment to the treatment of cancer, viruses, and microbial infections [1–4]. The battle against resistance has been largely fought at the molecular level, leading to an increasingly mature understanding of its underlying biochemical and genetic roots. At the same time, evolutionary biologists have long recognized resistance as a fundamentally stochastic process governed by the complex interplay between microbial ecology and evolutionary selection. The last decade, in particular, has seen a significant surge in efforts to develop and understand evolution-based treatment strategies to forestall resistance [5–16]. While the vast majority of this work focuses on spatially homogeneous environments, a number of recent studies, both theoretical and experimental, have demonstrated that spatial heterogeneity in drug concentration can dramatically alter the evolutionary dynamics leading to resistance [16–24]. On a practical level, the picture that emerges is somewhat bleak, as resistance evolution is dramatically accelerated in the presence of spatial gradients in drug concentration [18–20, 22–24] or heterogeneous drug penetration [17, 21]. Interestingly, however, recent work shows that this acceleration can be tempered or even reversed when the mutational pathway (i.e. the genotypic fitness landscape) leading to resistance contains fitness valleys [18], which are known to inhibit evolution [25–28]. Unfortunately, because the fitness landscape is a genetic property of the cells themselves, the potential for accelerated evolution appears to be “built in”, making it difficult to combat in a treatment setting. However, these results raise the question of whether non-monotonic profiles of tunable properties of the system—for example, the spatial selection pressure— might also have the potential to slow evolution, even when the mutational pathway lacks the requisite fitness valleys.

Evolution in natural or clinical settings takes place in heterogeneous environments characterized by spatial fluctuations in multiple factors, including drug concentrations, nutrients, temperature, and pH, all of which potentially affect cellular growth. Understanding evolution and ecology in such spatially extended systems is a challenging and long-studied problem [29–33]. Recent studies have demonstrated rich dynamics when inter-cellular interactions are defined on heterogeneous complex networks [34–36], where spatial structure can (for example) promote invasive strategies in tumor models [35] or modulate fixation times on random landscapes [34]. Remarkably, in the weak selection limit, evolutionary dynamics can be solved for any population structure [36], providing extensive insight into game-theoretic outcomes on complex networks. In addition, theoretical tools from statistical physics have proven useful for understanding spatiotemporal dynamics in spatially structured populations in a wide range of contexts, including biologically-inspired Monte Carlo models [18], toy models of source-sink dynamics [19], stepping-stone models of spatial pattern formation [37], models of dispersion [38–42], and Moran meta-population models [43–45]. In a similar spirit, here we use stochastic models of evolution along with theoretical tools from statistical physics to investigate the effects of spatially heterogeneous fitness pressures on the evolution of resistance. In contrast to previous models defined on heterogeneous networks at the single-cell level, here we consider meta-populations connected via simple topologies and investigate the effects of spatial structure imposed by arbitrary distributions of selection pressure. While several elegant approaches exist for studying these models in particular limits (e.g. with a center manifold reduction) [43–45], here we instead use a classical mean first passage time approach based on adjoint equations to reduce the calculation of mean fixation times to a simple collection of linear equations that can be easily solved for arbitrary spatial distributions of selection pressures. This method also allows us to find the fixation times from arbitrary initial states, which are often difficult to compute using other methods. Using this approach, we show that resistance evolution can be either accelerated or decelerated by spatial heterogeneities in selection pressure, even when the spatially averaged selection pressure remains constant. In addition, we demonstrate that tuning the spatial distribution of selection pressure can dramatically slow fixation when the subpopulations of resistant mutants are not uniformly distributed in space.

To investigate resistance evolution on a spatially heterogeneous landscape, we consider a stochastic Moran-like model [46] of a finite population (*N*) consisting of (*N* – *n*\*) wild-type cells with fitness *r*_0_ ≤ 1 and *n*\* drug-resistant mutants with fitness *r*\*, which we set to unity without loss of generality. Note that this model does not include a fitness cost to resistance (i.e. *r*\* ≥ *r*_0_ for all conditions). At each time step, cells are randomly selected for birth and death, with cells of higher fitness (in this case, resistant cells) chosen preferentially for division (see SI for full model with transition rates). Wild-type cells can mutate to become drug resistant at rate *μ*; we neglect reverse transitions to the drug-sensitive state. To incorporate spatial heterogeneity, we consider a simple spatially extended system with *M* distinct microhabitats, each containing *N* cells; cells are allowed to migrate at rate *β* between connected microhabitats (Fig. 1a). At each time t, the state of the system is characterized by the vector *n*\*(*x_i_*) whose components correspond to the number of mutants in each discrete microhabitat *x_i_* = 0,1,…, *M* – 1. The system evolves according to a continuous time master equation

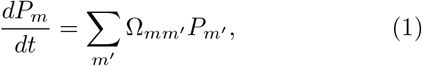

where *m* and *m*′ denote different states of the system and Ω is a *N^M^* × *N^M^* matrix whose entries depend on the wild-type fitness value *r*_0_(*x_i_*) at each spatial location *x_i_* (see SI). For tractability, we restrict our analysis to *M* = 3, which is the simplest model that allows for potentially non-monotonic fitness landscapes, such as fitness peaks and fitness valleys. In what follows, we refer to the vector *s*(*x_i_*) ≡ 1 – *r*_0_(*x_i_*) as the spatial profile of selection pressure, as it measures the difference in fitness between resistant and wild-type cells in each microhabitat (*x_i_*). Intuitively, large values of *s*(*x_i_*) correspond to regions where the resistant mutant has a significant evolutionary advantage over the wild-type cells (e.g. regions of high drug concentration).

**FIG. 1.**
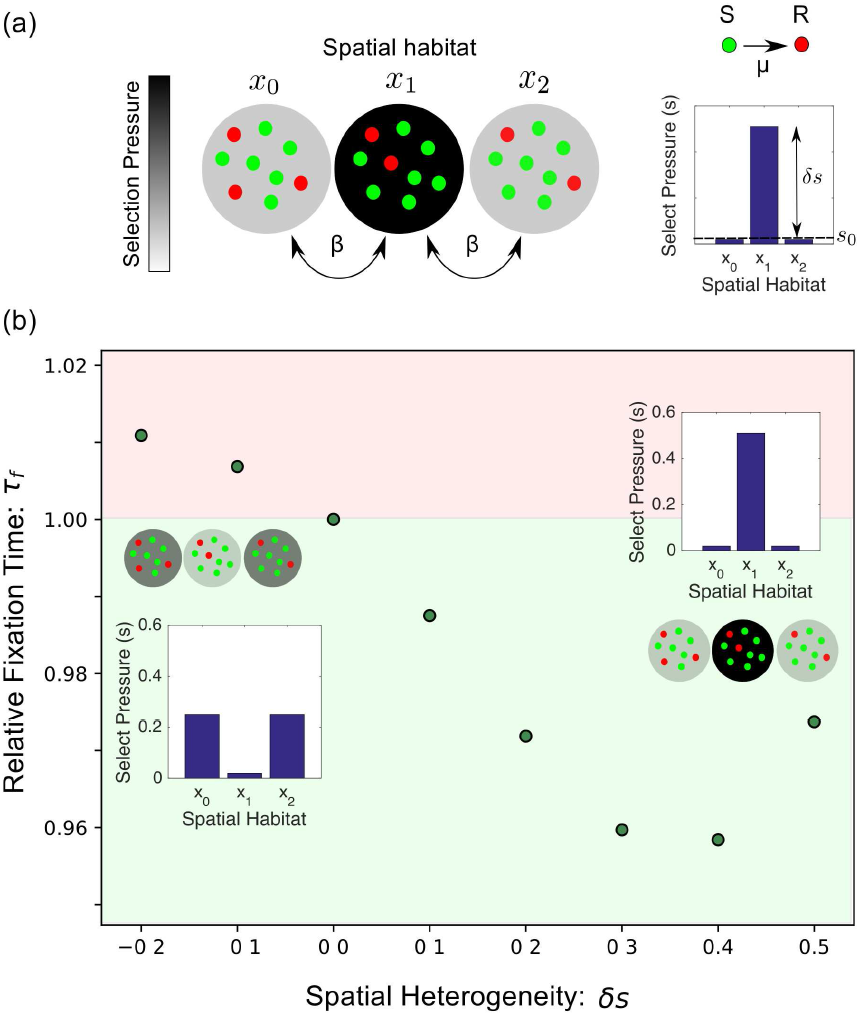
(a) Stochastic model for emergence and spread of resistant cells (red) in a spatially extended population of sensitive cells (green). Each spatial habitat (*x_i_*) contains *N* total cells. Cells migrate at a rate *β* between neighboring habitats, and sensitive cells mutate at a rate *μ* to resistant cells. The spatial distribution of selection pressure is characterized by a background value (*s*_0_) and a peak height (*δs*). (b) Example plot of the mean fixation time for different landscapes with *μ* = 5 × 10^−3^, *β* = 0.08, *N* = 25, and 〈*s*〉 = 0.167. The time to fixation can be either faster (green) or slower (red) than the spatially homogeneous landscape with *δs* = 0. Inset: selection landscapes for *δs* = −0.2 and *δs* = 0.5.

While Equation 1 is difficult to solve explicitly, it is straightforward to calculate quantities that describe the evolution of resistance in various spatial profiles. The model consists of a single absorbing state—the fully resistant state (*n*\*(*x_i_*) = *N* for all *x_i_*)—and the system will asymptotically approach this state. To characterize the speed of fixation in the presence of different spatial profiles *s*(*x_i_*), we calculate the mean first passage times (MFTPs) between states, which obey [47, 48]

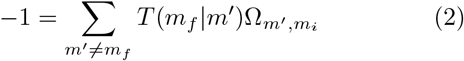

where *T*(*m_f_*|*m_i_*) is the mean time required for a system initially in state *m_i_* to first reach state *m_f_*. We take *m_f_* to be the fully resistant state and solve the coupled set of linear equations for 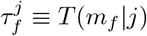, where *j* is an index that runs over all initial states. In particular, when *j* is the fully wild-type population (*n*\*(*x_i_*) = 0 for all *x_i_*), we refer to the MFPT as the mean fixation time *τ_f_*.

In the case of a single microhabitat, the mean fixation time *τ_f_* increases as selection pressure decreases (see SI). In the spatially extended case, *τ_f_* would also be expected to increase when the selection pressure is globally decreased, though it should also depend on the spatial structure of the specific selection profile *s*(*x_i_*). To investigate the impact of spatial structure alone, we compared *τ_f_* across different selection profiles *s*(*x_i_*), all of which were characterized by the same spatially averaged selection pressure, 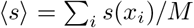. For simplicity, we begin with a symmetric profile characterized by a background selection pressure so in the edge habitats and a relative peak of height *δs* in the center habitat (Fig. 1a). This toy landscape has an average selection pressure of 〈*s*〉 = *s*_0_ + *δs*/*M*, and the parameters *s*_0_ and *δs* are constrained by the fact that 0 ≤ *s*(*x_i_*) ≤ 1 at all spatial locations. We vary *δs* systematically to explore different selection profiles, which can include a single selection pressure valley (*δs* < 0), a homogeneous landscape (*δs* = 0), or a single selection pressure peak (*δs* > 0).

Interestingly, we find that modulating heterogeneity (*δs*) can increase or decrease *τ_f_* for certain choices of migration and mutation rates, even when 〈*s*〉 is held constant (Fig. 1b). More generally, we find that the *β* – *μ* plane can be divided into three non-overlapping regions where the homogeneous landscape 1) leads to the smallest value of *τ_f_*, 2) leads to the largest value of *τ_f_*, or 3) does not correspond to an extremum *τ_f_* (Fig. 2a–b). In the latter region, heterogeneity often modulates the fixation time by only a few percent, but we do find larger effects in the high and low migration limits (i.e. on the edges) of the intermediate regime (Fig. S1). In addition, as we increase *β* for a fixed value of *μ, τ_f_* smoothly transitions from being minimized at *δs* = 0 to being maximized near *δs* = 0 (Fig. S1). We find empirically that the fixation time can be dominated by *τ*_1_, the time required to achieve a small population of mutants (Fig. 2c, rightmost panel) or *τ*_2_, the time required for this small population to achieve fixation (Fig. 2c, leftmost panel). However, in many cases—particularly those close to the intermediate region where fixation can be accelerated or slowed by heterogeneity—both timescales contribute to the dynamics. While we restrict ourselves primarily to *N* = 25, 〈*s*〉 = 1/6, and to symmetric landscapes, we find qualitatively similar results (i.e. 3 distinct regions) for other values of 〈*s*〉 (Fig. S2), *N* (Fig. S3), as well as for permuted selection profiles (Fig. S4), globally coupled profiles (Fig. S4), and monotonic (gradient) selection profiles (Fig. S5).

**FIG. 2.**
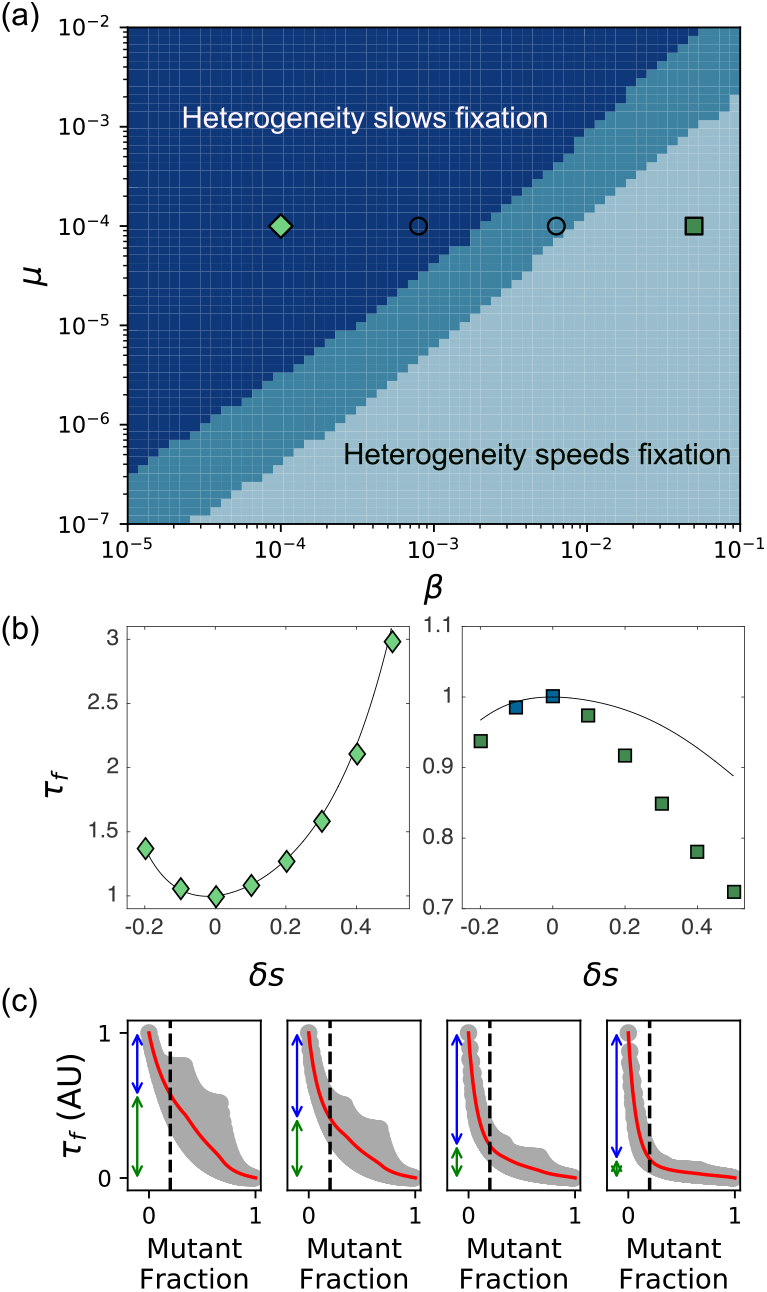
Spatial heterogeneity can speed or slow fixation depending on the rates of migration (*β*) and mutation (*μ*). (a) Phase diagram illustrates region of parameter space where the homogeneous landscape leads to a maximum (light blue), minimum (dark blue) or intermediate (medium blue) value of the in fixation time. MFPT calculations were performed for the indicated values of *β* and *μ* and for −0.2 ≤ *δs* ≤ 0.5 in steps of 0.1. (b) Sample fixation curves in the regions where heterogeneity slows fixation (left panel, diamonds; *β* = 10^−4^, *μ* = 10^−4^) or accelerates fixation (right panel, squares; *β* = 5 × 10^−2^, *μ* = 10^−4^). Solid curves indicate analytical approximations. (c) Gray shaded region indicates fixation time *τ_f_* from every initial state (*n*\*(*x*_0_),*n*\*(*x*_1_),*n*\*(*x*_2_)), where *n*\* (*x_i_*) is the initial number of mutants at position *x_i_*. Red curves show mean fixation time over all initial states with a given total mutant fraction. Vertical arrows represent time to achieve a total mutant fraction of 1/5 (*τ*_1_, blue) and time to go from that fraction to fixation (*τ*_2_, green). Left to right panels: increasing *β* at a fixed value of *μ* = 10^−4^; plots correspond to symbols on phase diagram in panel (a). *N* = 25 and 〈*s*〉 = 0.167 in all panels.

To intuitively understand these results, we developed a simple analytical approximation for *τ_f_* (see SI, Equation S16) valid in the limit *μ, β* ≪ 1, where the fixation time is dominated by the arrival times of individual mutants (either from de novo mutation or from migration from a neighboring vial that has achieved fixation). In this limit, the three habitats achieve fixation one at a time, and fixation in a single habitat is approximated as an exponential process with rate *λ*(*s, n_fix_*) = *N*(*μ + βn_fix_*)*P_fix_*(*s*), where *n_fix_* is the number of neighboring vials that have already achieved fixation and *P_fix_* = *s*(1 – (1 – *s*)^*N*^)^−1^ is the probability of a single mutant fixing in a habitat with selection pressure *s* (see SI). The approximation captures the qualitative features of fixation over a wide range of *μ* and *β* (Fig. S7) and, in many cases, provides excellent quantitative agreement as well (see, for example, Fig. 2b, left panel and Fig. S7).

In general, the analytical approximation for *τ_f_* is algebraically cumbersome. However, in the limit *β* ≪ *μ*, the approximation reduces to the expected maximum of three independent exponential random variables, leading to
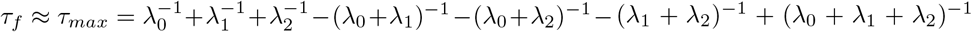, with *λ_i_* ≡ *λ*(*s*(*x_i_*), 0) (see SI for details). In this limit, the three-vial system acts effectively as three independent systems, with the overall fixation time corresponding to the slowest fixation. After rewriting *τ_max_* in terms of 〈*s*〉 and *δs*, it is straightforward to show that (∂*τ_max_*/∂*δs*)|_*δs*=0_ = 0 and (∂^2^*τ_max_*/∂*δs*^2^)|_*δs*=0_ > 0, indicating that the homogeneous landscape (*δs* = 0) minimizes the fixation time, consistent with results of the exact calculation (Fig. 2b, left panel). Intuitively, increasing heterogeneity reduces the minimum selection pressure in the spatial array, which in turn slows the expected maximum fixation time among the three habitats.

By contrast, in the limit *μ* ≪ *β, τ_f_* reduces to the expected minimum of three independent exponential processes, leading to *τ_f_* ≈ *τ_min_* = (λ_*eff*_)^−1^, where λ_*eff*_ ≡ λ_0_ + λ_1_ + λ_2_. In this limit, the fixation time is dominated by dynamics in the vial that first achieves fixation; the remaining vials then rapidly achieve fixation due to fast migration. For large but finite *N*, the fixation time *τ_min_* is maximized at *δs* = 0, indicating that heterogeneity always accelerates fixation, again consistent with the exact calculation (Fig. 2b, right panel). In this limit, the effective rate of fixation λ_*eff*_ is increased for all *δ* ≠ 0, as heterogeneity decreases fixation time in the vial with the fastest average fixation.

Our results indicate that a judicious choice of selection pressure profile can potentially slow fixation of de novo mutants. In addition, selection pressure profiles can be optimized to mitigate the effects of resistance once it has emerged. One advantage of the MFPT approach (i.e. solving Equation 2) is that it provides fixation times starting from all possible initial states, making it straightforward to apply to cases where a resistant subpopulation already exists. Specifically, consider a situation where a resistant subpopulation has arisen at a particular spatial location. Is it possible to choose the spatial distribution of selection pressure—for example, by spatially dosing the drug—to minimize the time to fixation from this state? Intuitively, the goal is to delay the onset of treatment failure as long as possible. As an illustrative example, we consider a population consisting of N/2 mutants in the center microhabitat and calculate the mean time to fixation for different spatial profiles of selection pressure. We then find the optimal value for δs—that is, the heterogeneity corresponding to the spatial landscape with the slowest fixation time—in different regions of parameter space (Fig. 3a). The specific choice of spatial profile significantly impacts the time to fixation from the initial resistant subpopulation (Fig. 3b). We observe two distinct regions of parameter space that lead to two very different dosing regimes (Fig. 3c). For *μ* sufficiently large relative to *β*, slowest fixation occurs when we maximize the amount of drug in the center microhabitat (*δs* = 0.5, white region). On the other hand, at large migration rates fixation is optimally slowed by maximizing the amount of drug in the two microhabitats without any initial mutants (*δs* = −0.2). In contrast to the case with no initial mutants (e.g. Figure 2), fixation time is never maximized by choosing the homogeneous profile. To further characterize these two regimes, we compare the fixation times from a maximally peaked landscape (*δs* is maximized) to that from a landscape with a large valley (*δs* is minimized). The selection landscape that leads to the slowest fixation rapidly becomes sub-optimal as mutation rate is decreased at constant *β* (Fig. 3d).

**FIG. 3.**
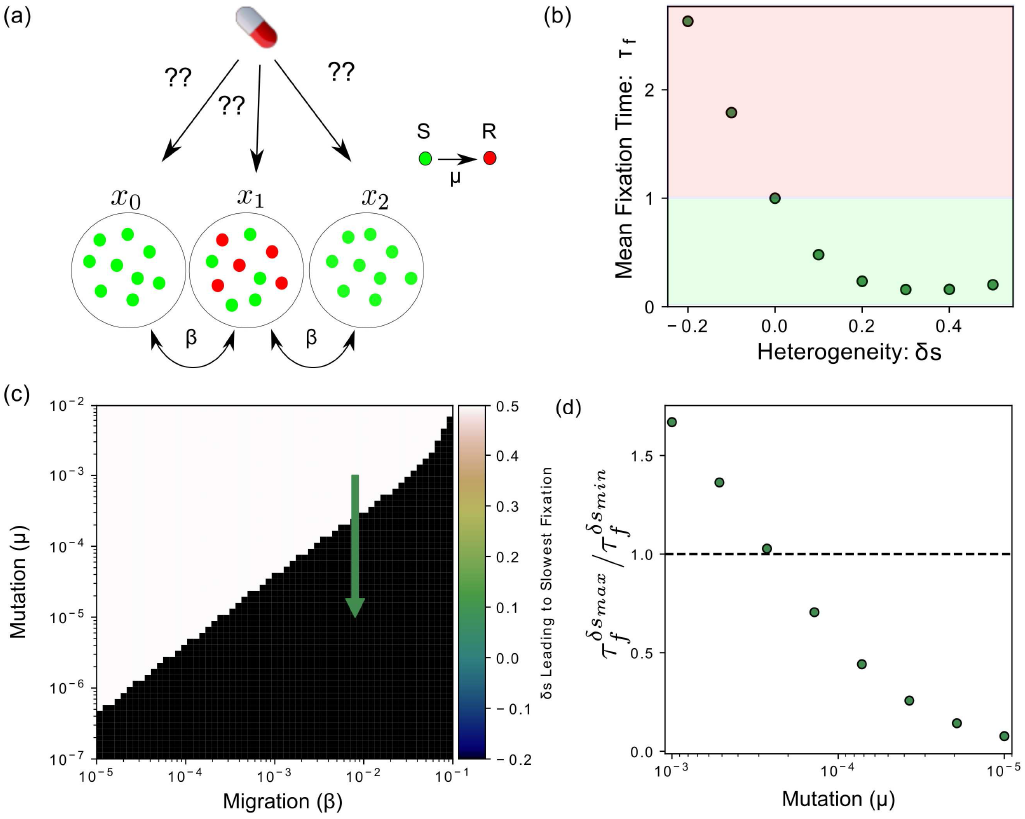
(a) Schematic: a subpopulation of resistant mutants (red) arises at a particular spatial location. How can one choose the spatial distribution of selection pressure (i.e. drug concentration) to maximize the time to fixation? (b) Heterogeneity can significantly speed or slow fixation starting from an initial resistant subpopulation consisting of *N*/2 cells in the center habitat (*μ* = 10^−5^, *β* = 8 × 10^−3^). (c) The optimal spatial heterogeneity (*δs*) leading to the slowest mean fixation time from an initial state of (0, *N*/2, 0). Depending on the specific parameter regime, the optimal selection pressure profile is the one with the largest possible valley consistent with 〈*s*〉 (black) or the one having the largest possible peak (white). (d) Relative magnitude of 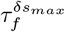 (mean fixation time at maximum value of *δs*) and 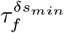 (mean fixation time at minimum value of *δs*) as mutation rate decreases at constant migration rate (green arrow, panel (c)). *N* = 25 and 〈*s*〉 = 0.167 in all panels.

Our model is a dramatic oversimplification of the biological dynamics leading to drug resistance. Practical applications will require analysis of more realistic models and may call for spatial optimizations with different constraints-for example, limits on the maximum allowable local selection pressure. Nevertheless, the simplicity of our model allows for a thorough characterization of fixation time over a wide range of parameters, and its behavior is surprising rich. Importantly, our results do not require a fitness cost of resistance or a genetic fitness valley, and they predict that spatial heterogeneity in drug concentrations would impact populations of motile and non-motile cells in opposing ways, even when mutations rates are relatively similar. While heterogeneity is likely to accelerate evolution for populations of motile bacteria, similar to what is observed in experiments with *E. coli* [22, 24], our results predict slowed evolution for less motile cells (e.g. the nosocomial pathogen *E. faecalis* [49]) or cells with rapid mutation rates. Perhaps most interestingly, our results suggest counter-intuitive, spatially optimal profiles for slowing the spread of resistance sub-populations. In the long term, these results may lay the groundwork for optimized, spatially-resolved drug dosing strategies for mitigating the effects of drug resistance.

